# Warming in the land of the midnight sun: breeding birds may suffer greater heat stress at high- vs low-Arctic sites

**DOI:** 10.1101/2021.10.26.465986

**Authors:** Ryan S. O’Connor, Audrey Le Pogam, Kevin G. Young, Oliver P. Love, Christopher J. Cox, Gabrielle Roy, Francis Robitaille, Kyle H. Elliott, Anna L. Hargreaves, Emily S. Choy, H. Grant Gilchrist, Dominique Berteaux, Andrew Tam, François Vézina

## Abstract

Rising global temperatures are expected to increase reproductive costs for wildlife as greater thermoregulatory demands interfere with essential breeding activities such as parental care. However, predicting the temperature threshold where reproductive performance is negatively impacted remains a significant hurdle. Using a novel thermoregulatory polygon approach, we predicted the threshold temperature at which an Arctic songbird–the snow bunting (*Plectrophenax nivalis*)–would need to reduce activity and perform below the 4-times basal metabolic rate (BMR) required to sustain nestling provisioning to avoid overheating. We then compared this threshold to operative temperatures recorded at high (82°N) and low (64°N) Arctic sites to estimate how heat constraints translate into site-specific impacts on sustained activity level. We predict buntings would become behaviourally constrained at operative temperatures above 11.7°C, whereupon they must reduce provisioning rates to maintain thermal balance. Low Arctic sites had larger fluctuations in solar radiation, producing consistent daily periods when operative temperatures exceeded 11.7°C. However, high-latitude birds faced entire, consecutive days where parents would not be able to sustain required provisioning rates. These data indicate that Arctic warming is likely already disrupting the breeding performance of cold-specialist birds, but also suggests counterintuitive and severe negative impacts of warming at high-latitude breeding locations.

## 1. Introduction

Animals frequently experience stages that demand significant increases in their sustained rate of energy expenditure (e.g., reproduction [1–3]). In an era of rapid climate change that is impacting species and ecosystems worldwide [4], understanding energetic limits and their causes is paramount for predicting whether organisms can respond to current rising global temperatures [5]. Historically, energetic limits among endotherms have either been attributed to intrinsic physiological factors (e.g., the digestive capacity of the gut to assimilate energy; central limitation hypothesis [6]) or constraints residing in the metabolic capacity of specific peripheral tissues (e.g., mammary glands or muscle tissue; peripheral limitation hypothesis [1]). Recently, Speakman and Król [7–9] proposed an alternative hypothesis, termed the heat dissipation limit (HDL) theory, which contends that the maximal rate of energy expenditure for an endothermic animal is limited by physiological factors governing heat dissipation capacity and the consequent avoidance of lethal body temperatures. Importantly, whereas the peripheral limitation hypothesis argues that energetic constraints may act on a range of tissues and organs (e.g., mammary glands, brown adipose tissue, or skeletal muscle), the HDL theory proposes a universal constraint in the form of heat dissipation and thereby provides a mechanistic link between an animal’s physiological capacity to maximize energy expenditure with the interplay between heat dissipation and ambient temperature.

Despite the significant conceptual gains that the HDL theory has provided in linking heat dissipation capacity with energetic expenditure, our ability to predict the ambient temperatures that will ultimately constrain an animal’s performance (i.e., sustained rate of energy expenditure) remains a major impediment to assessing species vulnerability to climate change [10]. Although several studies have reported threshold temperatures above which sustained activity and/or reproductive performance were compromised [10–14], these studies derived threshold values from post-hoc analyses on behavioral observations (e.g., provisioning rates) and are therefore not predictive by design. Recently, Rezende and Bacigalupe [15] proposed a predictive analytical tool - the thermoregulatory polygon - for estimating the dimensional space in which thermoregulation is possible given an animal’s combined rate of energy expenditure and the environmental temperatures it is operating within. Thermoregulatory polygons are built from commonly measured physiological variables (basal and maximal metabolic rate, and minimum and maximum thermal conductance) to delineate the boundaries in which heat production and dissipation are balanced [15]. Thus, thermoregulatory polygons can help estimate animal responses to further warming by integrating concepts of the HDL theory to predict the ambient temperatures over which endothermic animals can sustain activity and avoid lethal body temperatures. Surprisingly, despite their potential as a predictive tool, to our knowledge only one study has applied thermoregulatory polygons, using them to predict the energetic consequences of activity time in nocturnal and diurnal mammals [5].

Among endotherms, birds are expected to be particularly sensitive to increasing environmental temperatures [16,17]. The offspring-rearing period for parents with dependent young requires substantial increases in sustained work effort, with adults typically performing at 4 to 6 times their basal metabolic rate (i.e., resting rate of energy expenditure; [2,3,6]). Any excess heat generated as a by-product from foraging and provisioning must ultimately be dissipated, or birds risk overheating (hyperthermia). Problematically, birds often breed during the warmest parts of the year when it is hardest to passively shed body heat [18]. Indeed, birds often decrease activity on days with warmer ambient temperatures, which is likely a thermoregulatory response to avoid heat stress [19–23]. Recent studies have also shown that when a bird’s capacity to dissipate body heat is increased (e.g., by experimentally removing insulative feathers), provisioning adults can sustain higher levels of activity and invest more in both their current and future reproductive efforts [24–27]. Thus, reproductive performance can be constrained by a bird’s capacity to dissipate body heat produced during essential breeding activities, suggesting that increasing environmental temperatures could significantly impact reproductive investment.

Here, we apply a thermoregulatory polygon to snow buntings (*Plectrophenax nivalis*; figure 1*b*), an Arctic-breeding songbird, to investigate how environmental temperature affects the interaction between thermoregulation and sustained energy expenditure on the breeding grounds. Applying thermoregulatory polygons to Arctic endotherms is extremely pertinent and valuable for predicting how increasing ambient temperatures under climate change will impact essential life-history stages through thermal constraints on behavior. Many Arctic animals are cold specialists and regularly endure extremely cold weather and have evolved physiological adaptations for minimizing heat loss [28–31]. Consequently, high-latitude breeding species are likely vulnerable to moderate increases in ambient temperature [32–35]; an alarming fact given that the Arctic has warmed faster than the global average and is expected to continue outpacing the global average over the 21^st^ century [4]. In addition, O’Connor et al. [32] recently showed that buntings in particular can become heat-stressed at even moderate air temperatures and that their evaporative cooling capacity is extremely limited. Consequently, highly active, breeding snow buntings exposed to constant solar radiation and modest rises in air temperature would be largely dependent on behavioural thermoregulatory strategies (e.g., reducing provisioning effort) as opposed to physiological mechanisms (e.g., sustained increases in evaporate water loss rates) to dissipate body heat and avoid lethal body temperatures.

**Figure 1.**
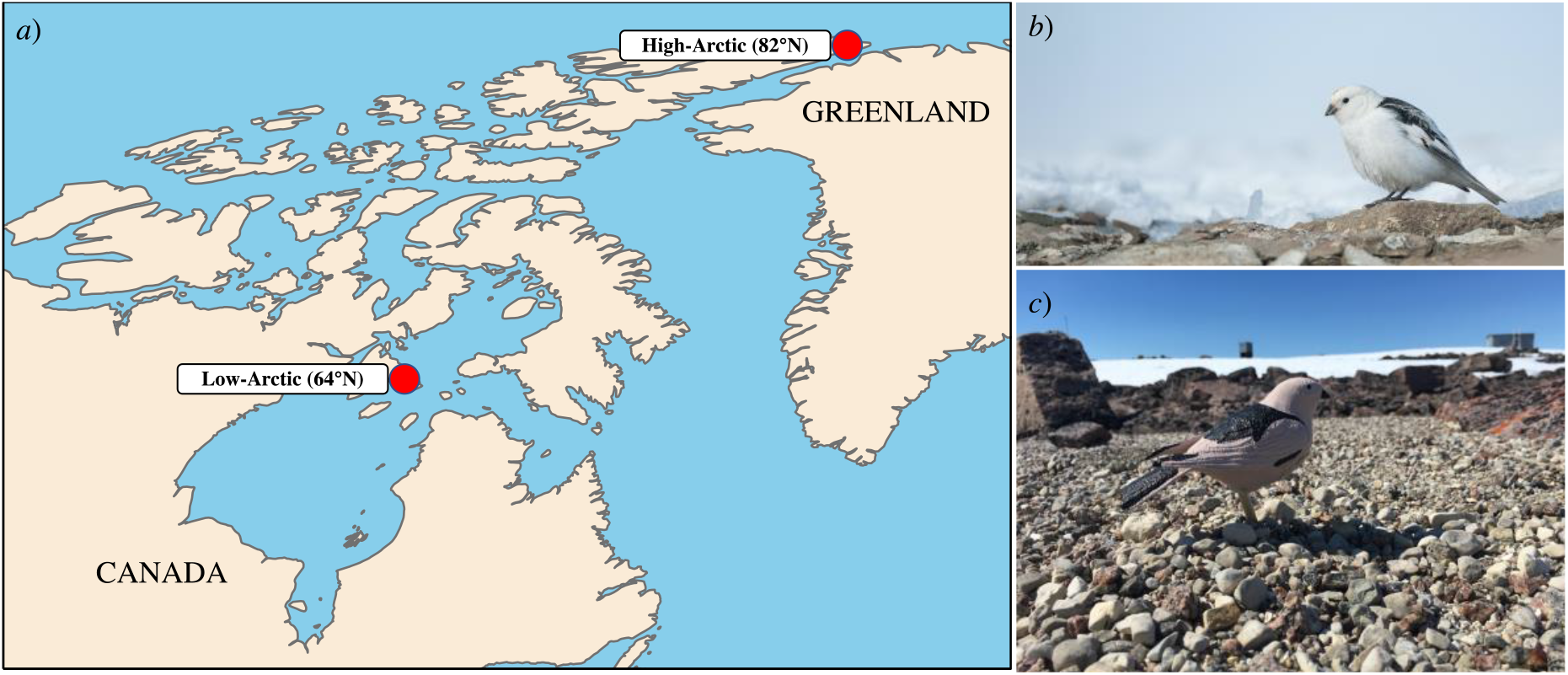
*a*) The location of the low-Arctic (East Bay Island, 64°N) and high-Arctic (Alert, 82°N) study sites. *b*) Photo of a snow bunting at the high-Arctic study site. *c*) Photo of a 3D printed snow bunting model in the field at the low-Arctic study site.

Our goal was to estimate how sensitive snow buntings’ performance may be to increasing Arctic temperatures, given their heat dissipation capacity. We first used thermal physiological data to construct a thermoregulatory polygon and predict the threshold temperatures at which sustainable performance (e.g., birds actively provisioning nestlings) would be expected to decline in buntings maintaining thermal balance (i.e., heat produced = heat dissipated). We then compared the thermoregulatory polygon prediction to both operative and air temperatures measured in the field at two latitudinally distant breeding sites to evaluate how heat constraints on bunting performance (i) differed between a low and high Arctic region, and (ii) could translate into site-specific impacts on reproductive performance and success.

## 2. Materials and methods

### (a) Operative and air temperature measurements

We measured operative and air temperatures during the snow bunting breeding period at two research sites in northern Canada representing the low-Arctic (East Bay Island; 64°01’N, 81°47’W; [36]) and high-Arctic (Alert; 82°30’N, 62°20’W; [37]; figure 1*a*). Operative temperature represents the temperature of the thermal environment as perceived by an individual and integrates the physical properties of the animal with the thermal properties of the local environment (e.g., air temperature, radiation, and wind; [38]). To measure the operative temperature perceived by snow buntings at our two sites, we used 3D printed, hollow plastic model birds (hereafter 3D models; [39,40]; figure 1*c*). We printed the 3D models to match the size and shape of an adult snow bunting (see electronic supplementary material, figure S1 in appendix S1). Additionally, we painted the 3D models to match the spectral properties of male snow buntings in breeding plumage, thereby allowing the 3D models to act as operative temperature thermometers. We painted the 3D models to match the color morph of male buntings because males feed females while incubating [41] and consequently are exposed to solar radiation for a longer duration throughout the breeding period. We also focused on males given their simplified monochromatic breeding plumage [42] and because males actively provision offspring at similar rates to females [43]. We used a spectrophotometer (Ocean Optics Jaz spectrometer) to measure the spectra of the black (N = 16 birds) and white (N = 27 birds) feather regions of male snow buntings within the 300-700nm wavelengths. We used the *pavo* package in R [44] to convert the spectra wavelengths to a red:green:blue (R:G:B) color combination. We then used an R:G:B-to-paint converter (https://www.e-paint.co.uk/convert-rgb.asp) to acquire a paint that best matched the R:G:B color combination of male bunting feathers. We opted to paint the 3D models (N = 68 at the high-Arctic site and N = 13 at the low-Arctic site) instead of placing the skin and plumage of a male snow bunting over the models as this optimized our experimental design by allowing us to record operative temperature in numerous models simultaneously across a broader geographic area [45].

We measured the internal temperature of each 3D model by placing a temperature logger in the centre of each model. At the high-Arctic site, we drilled a hole in the belly of each 3D model and secured an iButton (model DS1921G-F5, Maxim Integrated, San Jose, CA USA) in the approximate center (figures S2 and S3 in appendix S1) by gluing it to the end of a wooden dowel surrounded by a rubber stopper, creating an airtight seal around the drill-hole (figure S4 in appendix S1). At the low-Arctic site, models were similarly set up except for using Hobo data loggers (Pendant model, MX2201, Onset Inc., Bourne, MA USA) instead of iButtons, which we secured with silicone caulking. At both sites, the 3D printed models were secured to a wooden plank by gluing a wooden dowel to a notch in the 3D model (figures S3 and S4 in appendix S1). We cut the wooden dowels to approximate the height of a standing snow bunting. We covered each plank in the field using the substrate beneath the models to mimic the thermal properties of snow bunting’s natural environment (e.g., snow, moss, or rocky shale; figure S5 in appendix S1).

At each site, we deployed 3D models within representative breeding territories and across naturally occurring habitats to adequately capture the thermal heterogeneity experienced by buntings. In the high-Arctic, we recorded operative temperatures every 5 minutes from 22 May to 7 September 2019 and models were deployed over six separate periods, each lasting approximately 7 days (due to iButton memory limitations). After 7 days, we downloaded the operative temperature data and redeployed the 3D models to a new location. In the low-Arctic, we originally deployed 22 models once and recorded operative temperatures continuously from 11 June to 19 July 2019 at 2-min intervals. Unfortunately, polar bears damaged many models, so only 13 of our original 22 models were usable. However, the distribution of snow bunting breeding pairs at the low-Arctic site is much smaller in area than in the high-Arctic and these 13 models thus still provided sufficient geographic coverage of East Bay’s microclimates experienced by snow buntings.

At both study sites, we collected air temperature data to compare against operative temperatures. In the high-Arctic, the downloaded meteorological data was measured at the National Oceanic and Atmospheric Administration’s (NOAA) broadband radiation station located adjacent to the Global Atmospheric Watch (GAW) Observatory (82°28’N, 62°30’W). These data are 1-min averages of air temperature obtained at a height of 3 m above the ground using an aspirated Vaisala HMP-235 (PT100 sensor). In the low-Arctic, we collected an air temperature value every 30-min using six Kestrel weather meters (model 5500, Boothwyn, PA, USA) placed 2-3 m above ground level at separate locations across the study site.

### (b) Thermoregulatory polygon parameters and construction

Thermoregulatory polygons use an animals rate of energy expenditure and thermal conductance properties to delineate the space in which endotherms can balance heat production with heat loss under a given environmental temperature [15]. We calculated the basal metabolic rate (BMR; N = 28 birds), minimum wet thermal conductance (C_min_; N = 20 birds) and maximum dry thermal conductance (C_max_; N = 21 birds) on a wild population of snow buntings in the high-Arctic from 2 June to 25 July 2018. Information on gas analyzers, experimental protocol, body and air temperature measurements, and equations used for calculating metabolic rates are described in detail in Le Pogam et al. [37,46,47] and O’Connor et al. [32]. Briefly, we measured BMR overnight on fasted individuals resting inside a darkened metabolic chamber at thermoneutral temperatures (mean air temperature = 26.2 ± 0.8°C; note, air temperature inside metabolic chambers is equivalent to operative temperature [38]). For C_min_, we measured metabolic rates on individuals at a constant air temperature below their lower critical temperature of 10°C ([28]; mean C_min_ air temperature = -19.0 ± 1.8°C). We did not measure rates of evaporative water loss during our C_min_ runs and therefore for each bird we calculated minimum wet thermal conductance as follows:

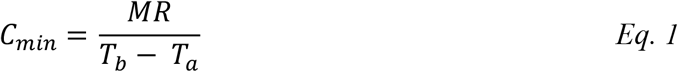

where MR represents metabolic rate in Watts, and T_b_ and T_a_ are the mean body and air temperatures, respectively. At air temperatures below the lower critical temperature, evaporative heat loss is minimal and thus its inclusion has little influence on C_min_ [48]. During metabolic measurements for C_min_, we measured T_b_ at the start and end of each run and used the mean value for our calculations.

We determined maximum dry thermal conductance by exposing birds to a ramped temperature protocol of gradually increasing air temperatures [32]. We only included birds that tolerated air temperatures above 31.5°C, representing the mean air temperature minus the standard deviation at which buntings started panting [32], as we assumed that birds that had initiated panting had reached their maximum thermal conductance [49]. This resulted in the removal of only 1 bird from the final data set. At higher air temperatures, evaporative heat loss becomes significant and therefore must be accounted for in the calculation of maximum thermal conductance [48]. We thus calculated maximum dry thermal conductance for each bird as follows:

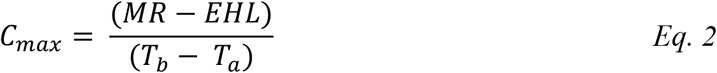

where EHL represents evaporative heat loss measured during respirometry trials [32]. During C_max_ experiments, we measured T_b_ continuously and therefore were able to calculate an average T_b_ over the same 5-min time window that metabolic rates were calculated [32].

To generate the thermoregulatory polygon, we calculated a combined mean across birds for each parameter (i.e, BMR, C_min_, C_max_ and T_b_). The BMR mean became the lower boundary of the thermoregulatory polygon. The C_min_ and C_max_ means became the slopes of the left and right boundaries, respectively. We calculated the y-intercepts for the C_min_ and C_max_ slopes using the equation:

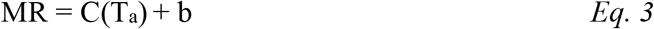

where C represents the combined C_min_ or C_max_ mean across birds and b is the y-intercept. We assumed T_a_ = T_b_ when MR = 0 [48] and used the combined T_b_ mean across birds during C_min_ (41.0 ± 0.4°C) and C_max_ (42.6 ± 0.7) measurements.

### (c) Estimating sustainable performance in the high-Arctic (Alert) and low-Arctic (East Bay)

We conducted all analyses in R 4.0.4 [50]. Over the course of the field season in the high-Arctic, we recorded a total of 843,773 individual operative temperature values from 68 3D models and a total of 107,092 air temperature values. In the low-Arctic, we recorded 405,000 individual operative temperature values from 13 models and a total of 10,803 air temperature values. We used these raw temperature data to create a time series of operative and air temperatures for each site averaged at 1-h intervals using the *timeAverage* function in the *openair* package [51].

The discontinuous sampling protocol in the high-Arctic (e.g., downloading data and redeploying models) resulted in 643 1-h gaps in our operative temperature time series. To estimate the percentage of time on a given day that buntings would have been behaviorally constrained from heat (see below) it was necessary to fill these gaps in our data. We filled the operative temperature gaps by fitting an artificial neural network [52–54] with the *neuralnet* package [55] to predict operative temperature based on seven radiative and meteorological variables observed at the NOAA broadband radiation station. Specifically, the input layer of the neural network included air temperature, wind speed (m/s), downwelling shortwave radiation flux (calculated as the sum of the contributions from diffuse and direct shortwave radiation; W/m^2^), reflected shortwave radiation flux (W/m^2^), albedo (calculated as the ratio of the reflected shortwave radiation flux to downwelling shortwave radiation flux), net longwave radiation flux (calculated by subtracting the longwave radiation flux emitted by the surface from the downwelling longwave radiation flux; W/m^2^), and diffuse fraction as a measure of the proportional influence of the direct sun and a proxy for cloudiness (calculated as the ratio of diffuse shortwave flux to total downwelling shortwave flux). Before training and testing the neural network, we applied a ranging standardization to the data, resulting in the data ranging between 0 and 1 [56]. We fitted the network with one hidden layer comprised of five neurons and trained the model using a random sample of 90% of the data set (1,752 values). The model was tested on a random sample of 10% of the data set (195 observations). We cross-validated the neural network by repeating the process (i.e., training, testing, and calculating mean square prediction error) 20 times consecutively. The neural network predicted hourly operative temperatures with an average mean square error of 1.8°C (range = 1.2 to 2.7°C).

We next used the C_max_ slope (i.e., the right-side upper boundary of the thermoregulatory polygon) to estimate the maximum sustainable energy expenditure of snow buntings maintaining thermal balance (i.e., heat production = heat loss) under both air temperature and operative temperature recordings. As the provisioning period is one of the most energetically expensive life-history stages for birds [3], we placed particular emphasis on the sustainable performance possible for buntings during this period. At the high-Arctic site, adult buntings are typically observed provisioning young from 4 July to 25 July (A. Le Pogam, personal observations) and at the low-Arctic site from 3 July to 24 July [42,43,57]. We thus used these respective date ranges to represent the typical provisioning period at each site. We defined performance as a multiple of BMR and assumed that 4-times BMR is the minimum sustainable performance required for adult buntings to adequately provision nestlings [2,3]. Therefore, we defined 4-times BMR as the energetic threshold for “optimal performance” and we calculated the percentage of time on a given day that buntings could work at either optimal (≥ 4-times BMR) or suboptimal (< 4-times BMR) performance levels based on either operative temperature or air temperature recordings. Lastly, we assumed buntings rested and significantly reduced provisioning rates for 3-h a day [58] and we only used temperature values measured between 01:00 and 22:00 hrs when calculating the daily percentage of time that buntings could work at optimal or suboptimal performance levels.

## 3. Results

### (a) Thermoregulatory polygon

All values reported are mean ± standard deviation. The mean basal metabolic rate (BMR) of snow buntings was 0.564 ± 0.076 W. Mean thermal conductance varied three-fold, with a calculated minimum wet thermal conductance of 0.023 ± 0.005 W/°C and a maximum dry thermal conductance of 0.073 ± 0.023 W/°C (figure 2*a*). The thermoregulatory polygon bounded by these physiological parameters predicts that snow buntings can maintain thermal balance and sustain optimal performance (i.e., ≥ 4-times BMR) at operative temperatures of up to 11.7°C (figure 2*b*). Once operative temperature exceeds 11.7°C, we expect buntings to become behaviorally constrained by heat and, consequently, forced to perform at suboptimal levels to avoid overheating. During the peak provisioning period when buntings are most active, operative temperatures over both sites ranged from -0.6°C to 24.6°C, leading to a 12.3°C zone in which we predict that buntings could maintain thermal balance and perform optimally (figure 2*b*).

**Figure 2.**
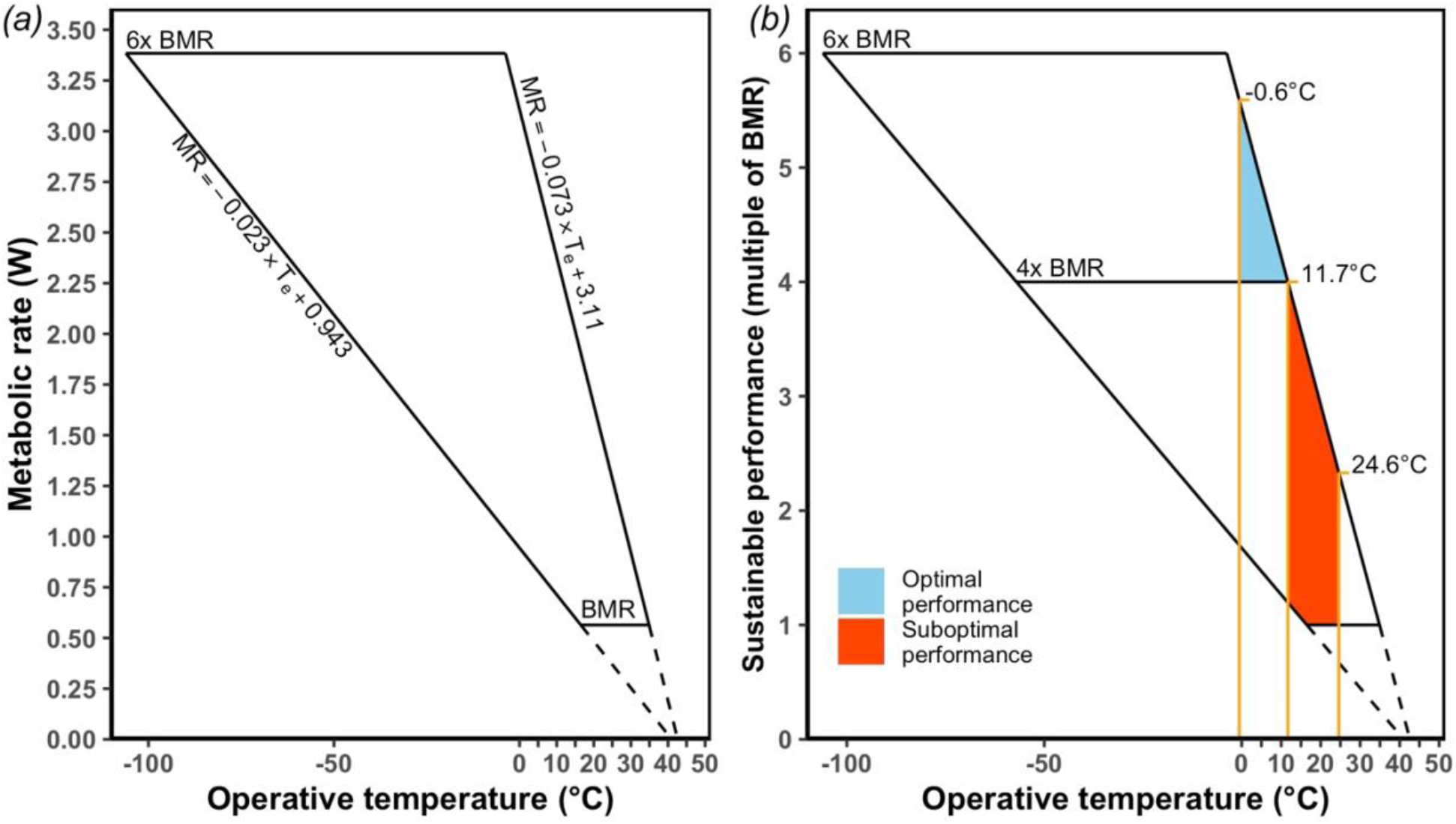
(*a*) Snow bunting thermoregulatory polygon bounded by basal metabolic rate (BMR; 0.564 W), minimum wet thermal conductance (0.023 W/°C), maximum dry thermal conductance (0.073 W/°C), and maximal sustained metabolic rate estimated at 6x BMR in breeding buntings. (*b*) Sustainable performance (expressed as a multiple of BMR) possible for buntings under thermal balance. At operative temperatures below 11.7°C, buntings can maintain thermal balance and sustain optimal performance (i.e., performance ≥ 4x BMR; blue zone). As operative temperatures increase, buntings must reduce activity, and concomitantly metabolic rate, to maintain thermal balance, resulting in a suboptimal performance (i.e., performance < 4x BMR; red zone). Optimal performance is defined as the sustained level of work required by adults to sufficiently rear nestlings. The black dashed lines are the extrapolation of the minimum and maximum thermal conductance slopes to the average body temperature recorded during laboratory measurements. The -0.6°C and 24.6°C temperatures on the right side of the polygon in *(b)* represent the average hourly operative temperature range measured in the field during snow buntings’ peak provisioning period.

### (b) Estimated sustainable performance in the high-Arctic (Alert) and low-Arctic (East Bay)

Air and operative temperatures at the high-Arctic site increased steadily from the beginning of the breeding period until peaking during the nestling-provisioning period, and then gradually declined towards the post-fledging period (see electronic supplementary material, figure S1*a* in appendix S2). Operative temperatures experienced by buntings frequently exceeded shaded air temperature, and on average were 3.5 ± 3.1°C warmer than air temperature (range of differences between operative and air temperature = -4.9°C to 14.5°C; figure S1*b* in appendix S2).

At the high-Arctic site, only operative temperature exceeded the predicted thermoregulatory polygon threshold value of 11.7°C before 5 July (figure 3*a*). However, from 5 July to 5 August, both air temperature and operative temperature measurements periodically exceeded 11.7°C (figure 3*a*), suggesting that buntings would have had to regularly perform at suboptimal levels below 4-times BMR during this period. Within the typical nestling-provisioning period at the high-latitude site (i.e., 4 July to 25 July) when buntings are most energetically active, modelling based on the polygon predicts that buntings experienced multi-day periods where they could have either performed at optimal levels for their entire active period (i.e., 01:00 hrs to 22:00 hrs) or they would have been heat constrained and forced to work below 4-times BMR for the entire active period (figure 4*a*). For example, based on operative temperature recordings, there were two periods of consecutive days (9 to 11 July and 19 – 22 July) where we predict that buntings would never have been heat constrained and could have worked at optimal performance levels for their entire active period (figure 4*a*). However, there were two periods of consecutive days (6 to 8 July and 13 to 17 July) when operative temperatures exceeded 11.7°C for their entire active period, and we predict that buntings would have to reduce their provisioning rates to lower metabolic heat production and avoid lethal body temperatures. From 13 to 19 July the polygon model also predicted that buntings experienced only 5 hours with temperatures that allowed them to both maintain thermal balance and sustain an optimal performance level of ≥ 4-times BMR. These findings suggest that temporal variation in heat constraints on the sustainable performance of breeding snow buntings in the high-Arctic correspond to synoptic cycles (i.e., weather-scale, 2-4 days) rather than regular circadian cycles. This is explainable by the suppressed amplitude of the diurnal cycle at the high latitude site where the sun is above the horizon continuously from early April through early September. Overall, under operative temperature recordings at the high-Arctic site, the percentage of time each day that buntings would have been behaviorally constrained from heat during their active period ranged from a minimum of 19% (4 hrs) to a maximum of 100% (21 hrs; figure 4*a*).

**Figure 3.**
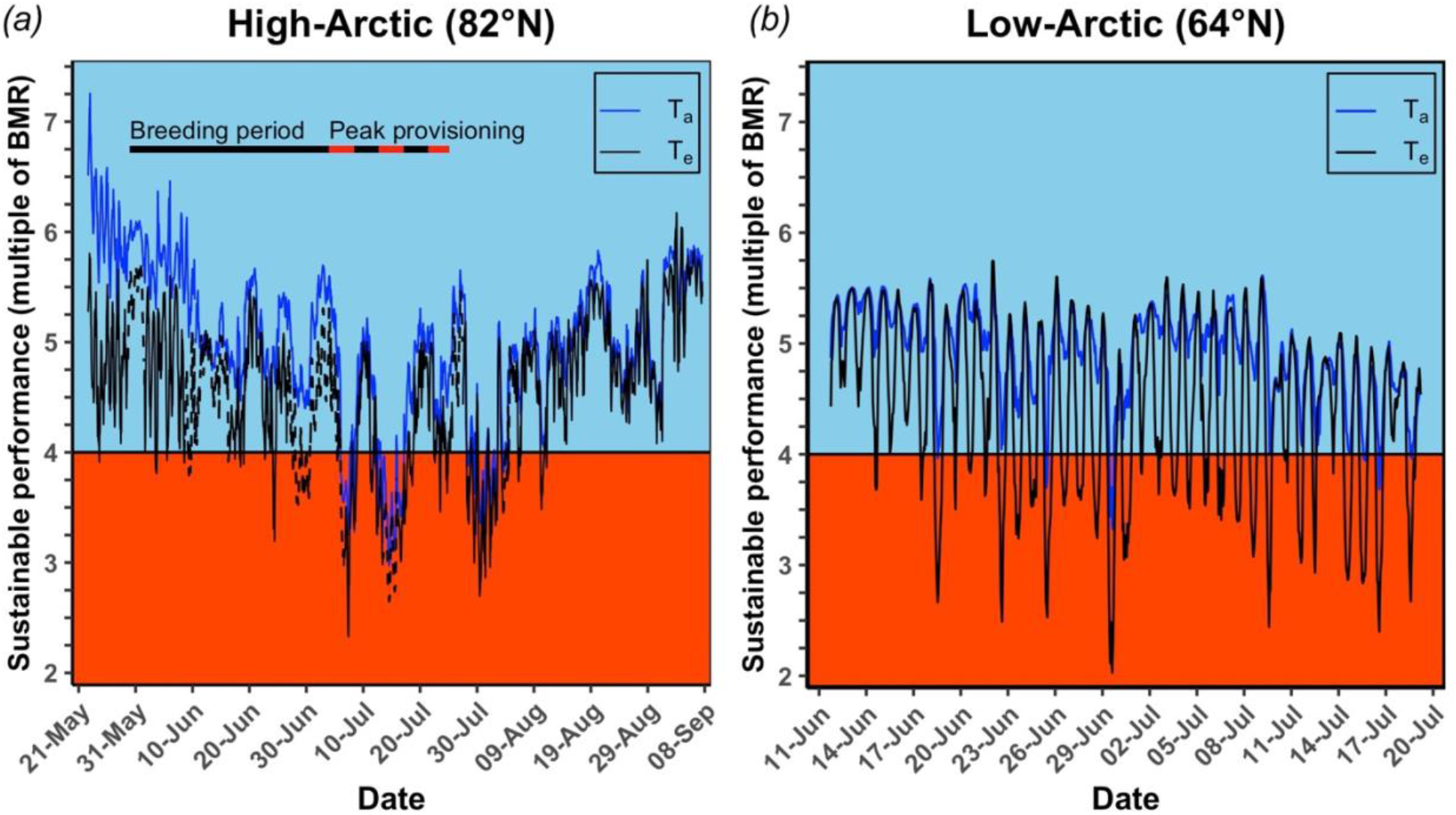
Estimated sustainable performance possible for snow buntings maintaining thermal balance at a (*a*) high-Arctic (Alert, Nunavut Canada) and (*b*) low-Arctic (East Bay Island, Nunavut Canada) breeding site. The line dividing blue and red zones represents the 4-times basal metabolic rate (BMR) minimum required for adults to maintain adequate provisioning to growing nestlings (see methods for details). The blue zone represents the times when average hourly operative (T_e_) or air (T_a_) temperature was below the thermoregulatory polygon threshold temperature of 11.7°C, predicting that buntings could sustain performance levels ≥ 4-times basal metabolic rate without altering behaviour. The red zone represents the times when T_e_ or T_a_ exceeded 11.7°C, predicting that buntings would be required to reduce their provisioning behaviour to subsequently work below 4-times basal metabolic rate to limit heat production and avoid lethal body temperatures. The dashed black lines in panel *a* represent the predicted T_e_ values from the artificial neural network (see methods for details).

**Figure 4.**
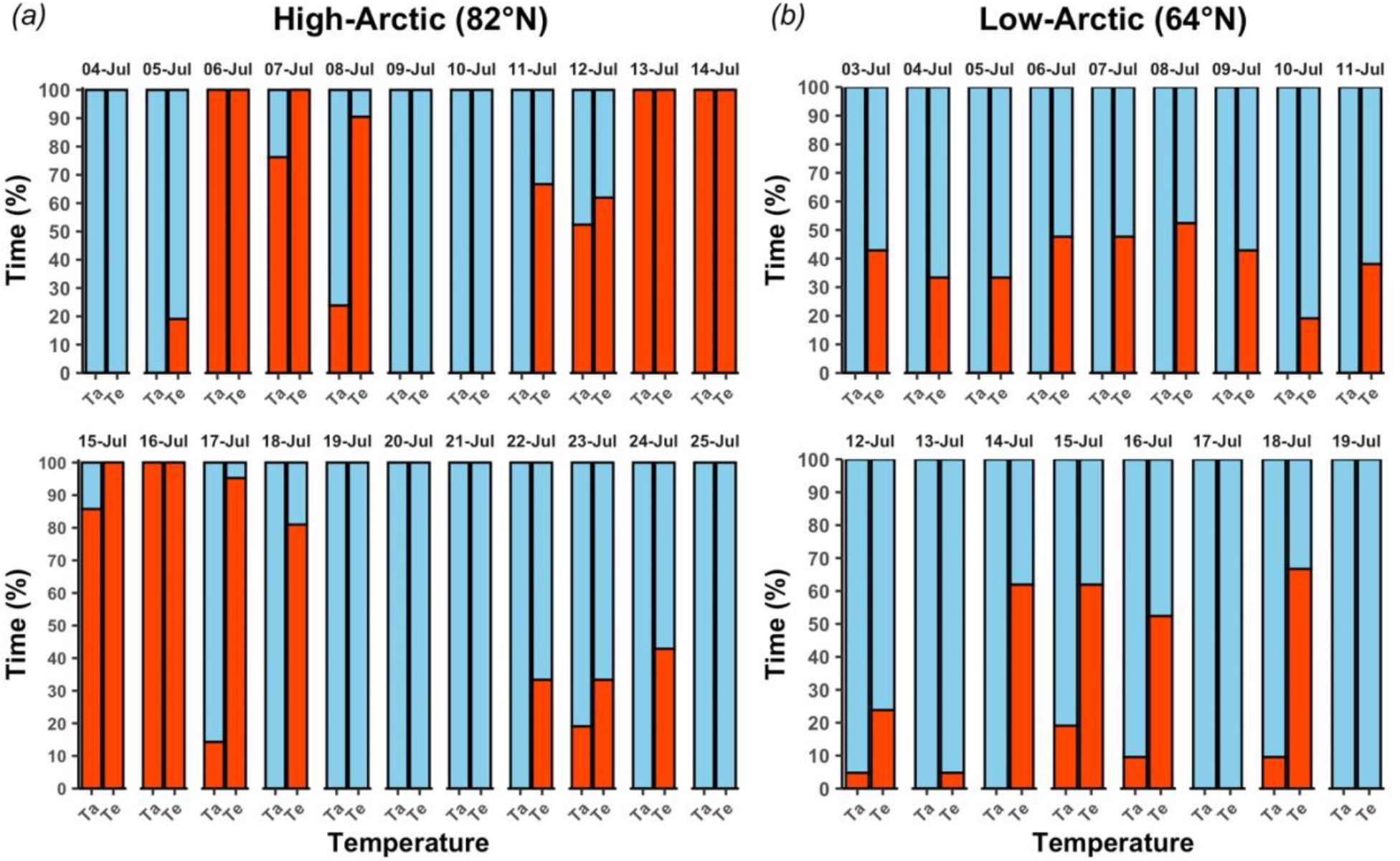
The daily percentage of time during their active period (01:00 hrs to 22:00 hrs) when snow buntings at a (*a*) high-Arctic (Alert, Nunavut Canada) and (*b*) low-Arctic (East Bay Island, Nunavut Canada) breeding site could either sustain an optimal performance level (blue region) or forced to work at suboptimal performance levels (red zones) based on either operative (T_e_) or air (T_a_) temperature recordings. Optimal and suboptimal performance is defined as the periods when buntings could sustain levels of work ≥ 4-times basal metabolic rate or < 4-times basal metabolic rate, respectively, while maintaining thermal balance under a given heat load.

At the low-Arctic site, average hourly temperatures were relatively consistent across the study period (figure S2*a* in appendix S2). The overall mean difference between operative and air temperature was 4.0 ± 4.1 (range = -2.7°C to 15.5°C; figure S2*b* in appendix S2). In contrast to the high-Arctic site, where both operative and air temperature recordings exceeded the thermoregulatory polygon threshold temperature of 11.7°C, only operative temperature recordings at the low-Arctic site consistently placed a heat constraint on buntings’ sustainable performance (figure 3*b*). For example, during the typical nestling-provisioning period in the low-Arctic (i.e., 3 July to 24 July), modelling based on the polygon predicted that buntings would have been behaviorally constrained on just four days using measured air temperature values, whereas operative temperature values suggest that bunting performance would have been constrained to some degree on 15 of the 17 days during which temperatures were recorded (figure 4*b*). Furthermore, unlike the high-Arctic birds, buntings at the low-Arctic site would not be forced to perform at suboptimal levels for their entire active period (i.e., an entire day), but would instead be forced to alter performance for a portion of each day (figure 4*b*). Overall, under measured operative temperatures at the low-Arctic site, the percentage of time that buntings would have been behaviorally constrained from heat on a given day during their active period (i.e., 01:00 to 22:00 hrs) ranged from a minimum of 5% (1 hr) to a maximum of 67% (14 hrs; figure 4*b*).

## 4. Discussion

A growing body of evidence suggests that increasing environmental temperatures associated with climate change will impose reproductive costs on birds via trade-offs between essential breeding behaviours and the need to dissipate body heat and avoid lethal body temperatures [10,11,17,25,59,60]. To date, predicting the threshold temperatures that will adversely affect breeding activity has been a limiting factor in forecasting the impacts of anthropogenic climate change on birds. Additionally, studies on how thermoregulatory demands will negatively impact breeding behaviour within birds is overwhelmingly biased towards hot, arid climates while studies on Arctic birds are severely lacking.

Using a thermoregulatory polygon approach, we estimated the maximal sustained energy expenditure in an Arctic songbird maintaining thermal balance across a range of environmental temperatures. Assuming an optimal performance level of 4-times basal metabolic rate (BMR; [3]), our findings predict that buntings will become heat constrained at operative temperatures above 11.7°C. At this point, buntings would need to reduce their maximal sustained energy expenditure and provision their offspring at sub-optimal performance levels to balance heat loads and avoid a lethal body temperature. Additionally, by examining impacts at both a low and high Arctic breeding site, our data reveal latitude-dependent operative temperature traces, likely linked to differences in available sunlight and solar radiation input, culminating in site-specific patterns in the heat constraints placed on an animal’s maximal sustained energy expenditure. Consequently, synoptic-scale influences on local temperature apparently dominate in modulating operative temperatures in the high-Arctic, whereas the diurnal cycle is the dominant factor in the low-Arctic. Collectively our results indicate that while Arctic warming will expose all snow bunting populations to more periods above their threshold temperature for sustained optimal performance, high-Arctic birds will likely face increases in the duration and magnitude of these periods due to the added effects of a higher average solar radiative flux. The expectation then will be that high-Arctic populations will face greater downstream costs to reproductive performance (e.g., provisioning rates), and therefore ultimately breeding success compared to low-Arctic populations.

### (a) Using the thermoregulatory polygon to predict thermal constraints

The heat dissipation limit theory postulates that an animal’s maximum sustained energy expenditure scales with its capacity to dissipate body heat [8]. Many factors influence an animal’s thermoregulatory ability, including basal metabolic rate and thermal conductance [61– 64]. Based on buntings’ basal metabolic rate (0.564 W) and maximum dry thermal conductance (0.073 W/°C), the thermoregulatory polygon predicts that at operative temperatures above 11.7°C, snow buntings cannot maintain thermal balance and sustain activity at optimal expenditure rates of 4-times basal metabolic rate. Therefore, if operative temperature exceeds 11.7°C for extended periods, we would expect to observe (i) a decline in the body condition of breeding adults as they increase evaporative water loss rates to maintain foraging effort and provisioning rates outside of the thermoregulatory polygon limits [11,59,65] and/or (ii) a slower growth rate, prolonged breeding period and potentially reduced fledging mass as adults reduce provisioning rates to avoid lethal body temperatures [13,41,66–68]. Although not derived under the thermoregulatory polygon framework, Cunningham et al. [12] reported lower provisioning rates at higher ambient temperatures in common fiscals (*Lanius collaris*) and fledglings were significantly lighter if maximum air temperature often exceeded a threshold temperature of 33°C. The comparatively low threshold temperature for buntings (11.7°C) likely stems from their physiological adaptions for life in the cold [37]. Consequently, snow buntings’ cold specialization appears to come at the cost of not being able to adequately dissipate heat through increases in maximum thermal conductance at even moderate operative temperatures.

Because the thermoregulatory polygon boundaries are set by the thermal conductance of the animal, they represent the thermal space in which an animal can balance heat loss and gain through sensible heat flow (i.e., non-evaporative mechanisms). Theoretically, an animal could maintain thermal balance and sustain a high rate of energy expenditure outside its thermoregulatory polygon by continuously dissipating body heat evaporatively. However, long-term evaporative water loss is not sustainable and evaporative cooling capacities vary significantly among species [16,69,70]. Recently, O’Connor et al. [32] showed that the evaporative cooling capacity of buntings is extremely limited, with most birds unable to evaporatively shed an amount of heat equivalent to their metabolic heat production. Therefore, it is unlikely that snow buntings can rely on evaporative cooling for prolonged periods to sustain activity outside their thermoregulatory polygon limits, and, instead, will be highly dependent on behavioral thermoregulation.

### (b) Site-specific impacts of thermal constraints on breeding performance and success

Solar radiation is a major driving force of operative temperature and can vary by time of day, year, or geographic location [52,71]. Our two sites represent the general southern and northern breeding limits for Arctic-breeding snow bunting populations in Canada [58], and are separated by ∼18° latitude. This latitudinal gap results in different quantities of solar radiation reaching the earth’s surface [52], likely producing the significant differences observed in the duration and frequency that operative temperature exceeded the predicted threshold temperature. For example, during the peak nestling-provisioning period, buntings at the high-Arctic site were predicted to frequently experience consecutive days where they would not be able to perform at 4-times their BMR. In contrast, buntings in the low-Arctic were predicted to experience shorter, but more consistent heat constraints on provisioning activity almost every day. Given that snow bunting nestlings have some of the highest recorded growth rates of any passerine (11.6-13.0 %/day of adult body mass; [72]), these latitudinal differences in constraints suggest that warming will produce different impacts on provisioning behaviour and hence offspring growth and survival at different populations. For instance, lower-latitude breeding birds could possibly make up for reduced provisioning opportunities each day by adjusting their activity budget throughout the day; working harder during the cooler periods to counteract overheating risks during warmer periods [5,73]. Indeed, under identical heat loads, Tapper et al. [27] observed higher feeding rates in wild female tree swallows (*Tachycineta bicolor*) that had their ventral feathers clipped to experimentally increase heat dissipation rates relative to unclipped females. Alternatively, parents breeding at lower latitudes could provision growing nestlings at lower rates per day and possibly extend the developmental period of the growing young. However, this could nonetheless impose survival constraints on nestlings and fledglings given that ground-nesting songbird species have evolved rapid growth rates and shorter in-nest development periods due to high rates of nest predation [74,75], as well as the short, ephemeral nature of productivity in insects required for offspring growth [76–78].

For higher latitude populations, the accumulation of reduced provisioning opportunities for consecutive days could impose substantial developmental costs on nestling development that may simply be too great for parents to compensate for on cooler days. Chick provisioning in buntings typically lasts 13 days; lowering provisioning rates for 3-4 consecutive days could have major impacts on chick condition at fledging, and possibly, post-fledging survival [60,79,80]. Consequently, as rapid Arctic warming continues [4], the temperature-dependent costs on reproductive performance may be more strongly felt at higher latitudes where climatic and meteorological patterns subject individuals to unique operative temperature cycles, with above threshold temperatures potentially lasting for days at the peak of breeding activities. It is worth noting, however, that our 3D models were painted to match the male color morph and therefore represent operative temperatures perceived by male snow buntings. During the provisioning period, both male and female buntings feed young and thus the operative temperatures experienced by females may differ from males leading to different sex constraints on performance. For instance, females lack the full dark back of male buntings and hence may experience lower operative temperatures allowing them to maintain higher provisioning rates than males. Nevertheless, under such a scenario, we would still predict negative impacts on nestling condition and fledgling success as both parents cannot adequately feed young at optimal rates.

The thermal environment experienced by wild animals represents a complex integration of biotic and abiotic factors operating at spatial scales relevant to the size of the animal [38,81,82]. The thermal environment, as measured by operative or standard operative temperature [83], can significantly deviate from air temperature [32,84]. Indeed, we found that operative temperature was, on average, 3.5 to 4.0 °C above air temperature but could exceed it by as much as 14.5 or 15.5°C. More importantly, our data show that the duration and frequency that the threshold temperature was exceeded markedly differed depending on the heat index used (i.e., air temperature or operative temperature). The implication of these differences is significant because any derived predictions for maximal sustained energy expenditure under increasing global temperatures will be substantially different whether air temperature or operative temperature measurements are used. For example, using air temperature alone at the low-Arctic site, we would predict that snow buntings are seldom heat constrained and can currently sustain optimal energy expenditures, despite operative temperatures indicating otherwise. In contrast, at the high-Arctic site during the peak provisioning period, air temperature measurements often exceeded the temperature threshold alongside operative temperature but the percentage of time that buntings were constrained was greater under operative temperature. Consequently, using air temperature alone, especially values derived at macroscales (e.g., WorldClim dataset at 1 km^2^ [85]), will certainly misrepresent an animals realized thermal environment, which operates at microscales, leading to biased and erroneous predictions on the impacts of climate change [86,87].

## Supporting information

3D model construction

operative and air temperature traces

## Acknowledgements

We sincerely thank the 2019 East Bay and Alert field teams for their assistance. We are grateful to Dr. Christina Semeniuk for the use of spectrophotometer measurement equipment and to Kevyn Gammie-Janisse and Chris Harris for their assistance with spectrophotometer measurements and analysis. We are also indebted to Lincoln Savi for providing essential help with the 3D models.

