## Supplementary material for "Warming in the land of the midnight sun: breeding birds may suffer greater heat stress at high- vs low-Arctic sites": 3D model construction

**Electronic supplementary material**

Construction of the 3D printed operative temperature thermometers

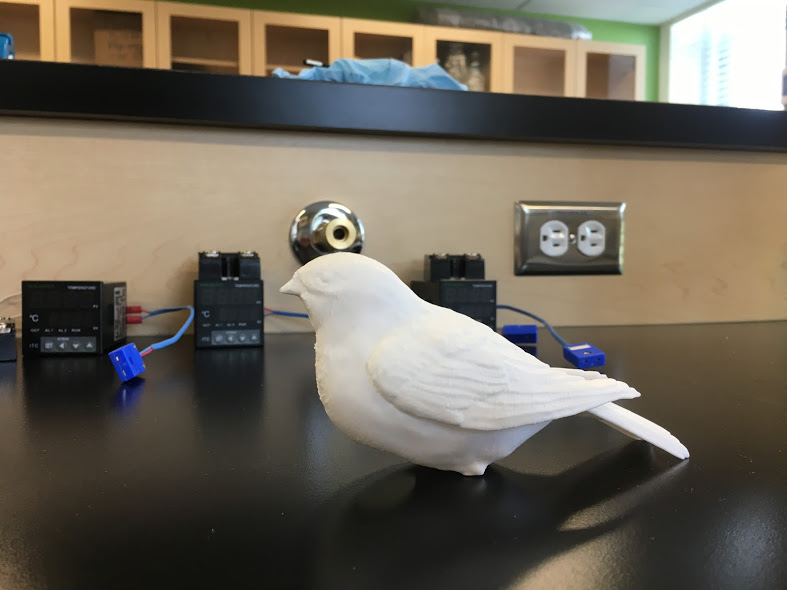

**Figure S1.** The finished 3D printed operative temperature thermometer before painting and temperature logger placement.

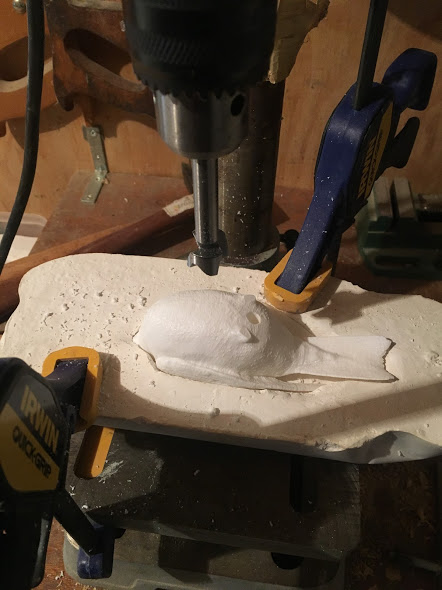

**Figure S2.** The placement of the 3D printed operative temperature thermometers in a plaster-of-paris mold used to secure the plastic bird when drilling the hole for the temperature logger.

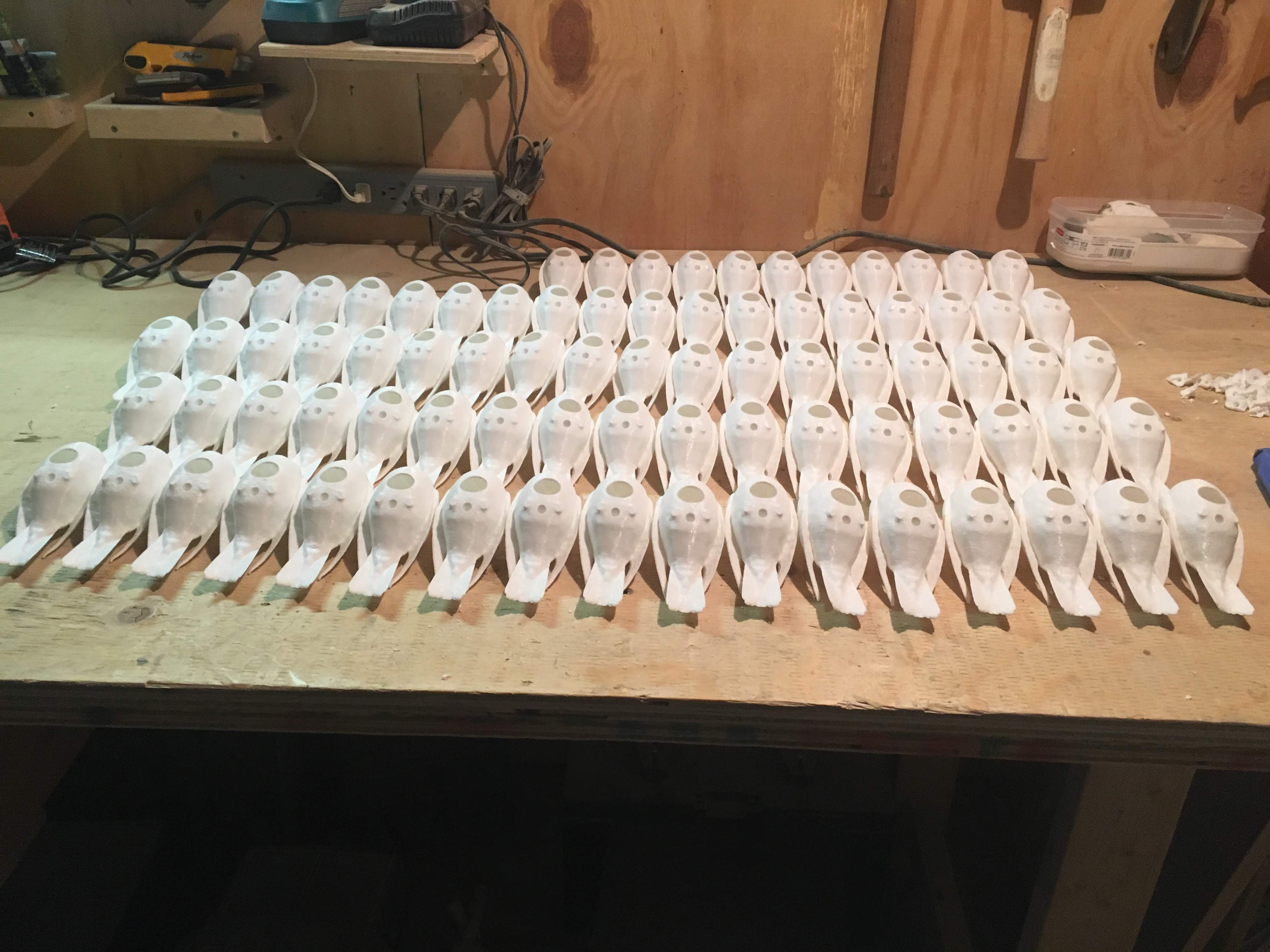

**Figure S3.** A Picture showing some of the drilled 3D printed birds before painting. The blue arrow is pointing to the drill hole for temperature loggers and the green arrow is the notch for the dowel which was secured to a wooden plank.

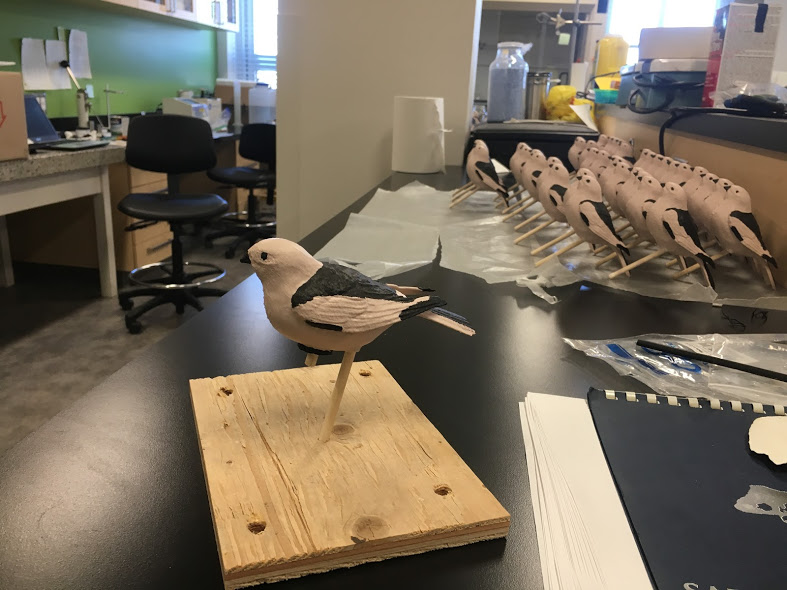

**Figure S4.** Picture showing the placement of the 3D printed operative temperature thermometer on the wooden plank. The blue arrow is pointing at the wooden dowel to which the temperature logger is glued and the rubber stopper that is plugging the drill hole. The green arrow is pointing at the wooden dowel which is placed in the notch indicated by the green arrow in Figure S3 above.

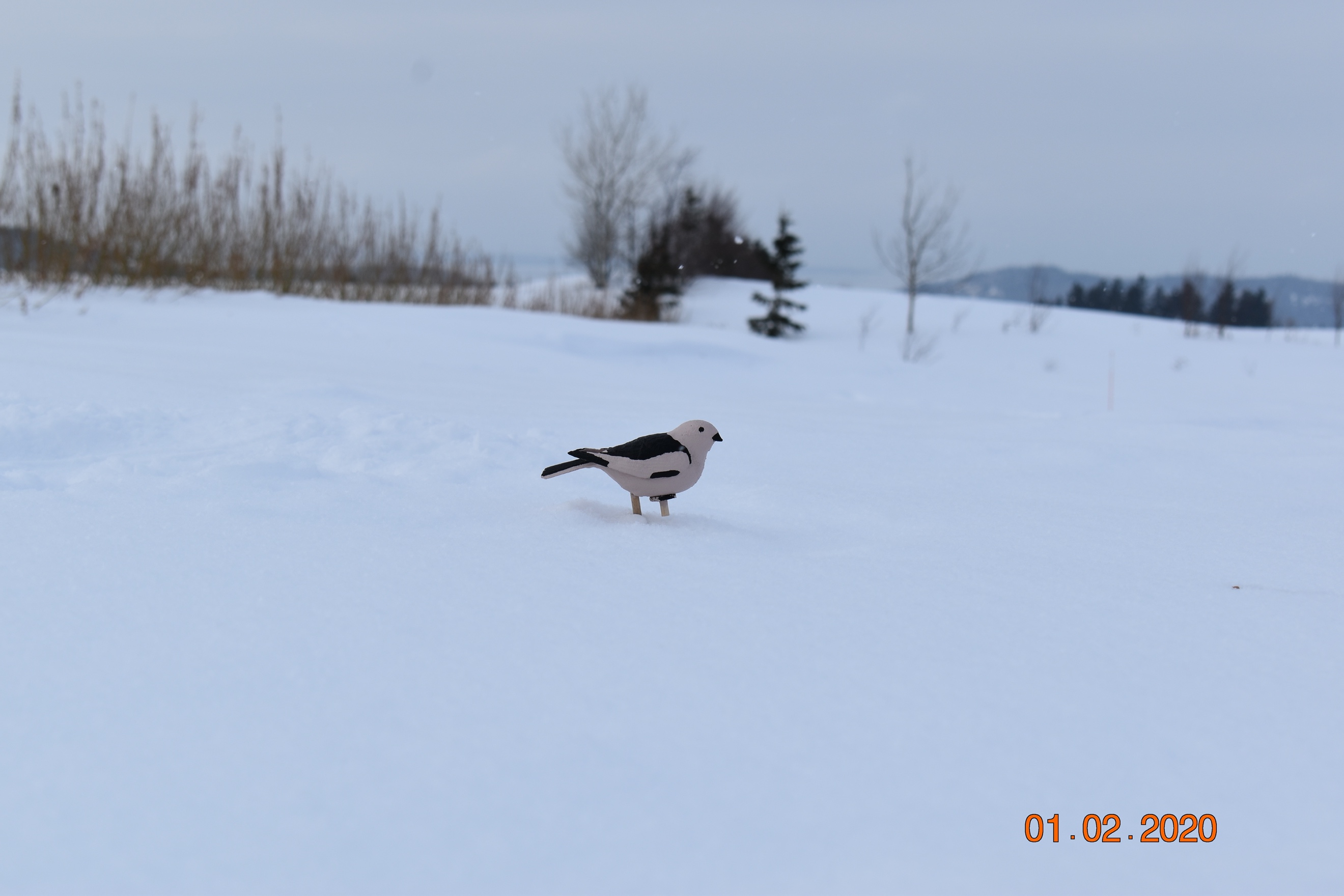

**Figure S5.** 3D model of a snow bunting placed in the field with snow placed over the wooden plank.
