## Supplementary material for "Warming in the land of the midnight sun: breeding birds may suffer greater heat stress at high- vs low-Arctic sites": operative and air temperature traces

**Electronic supplementary material**

1-hour averaged traces of operative and air temperature measurements

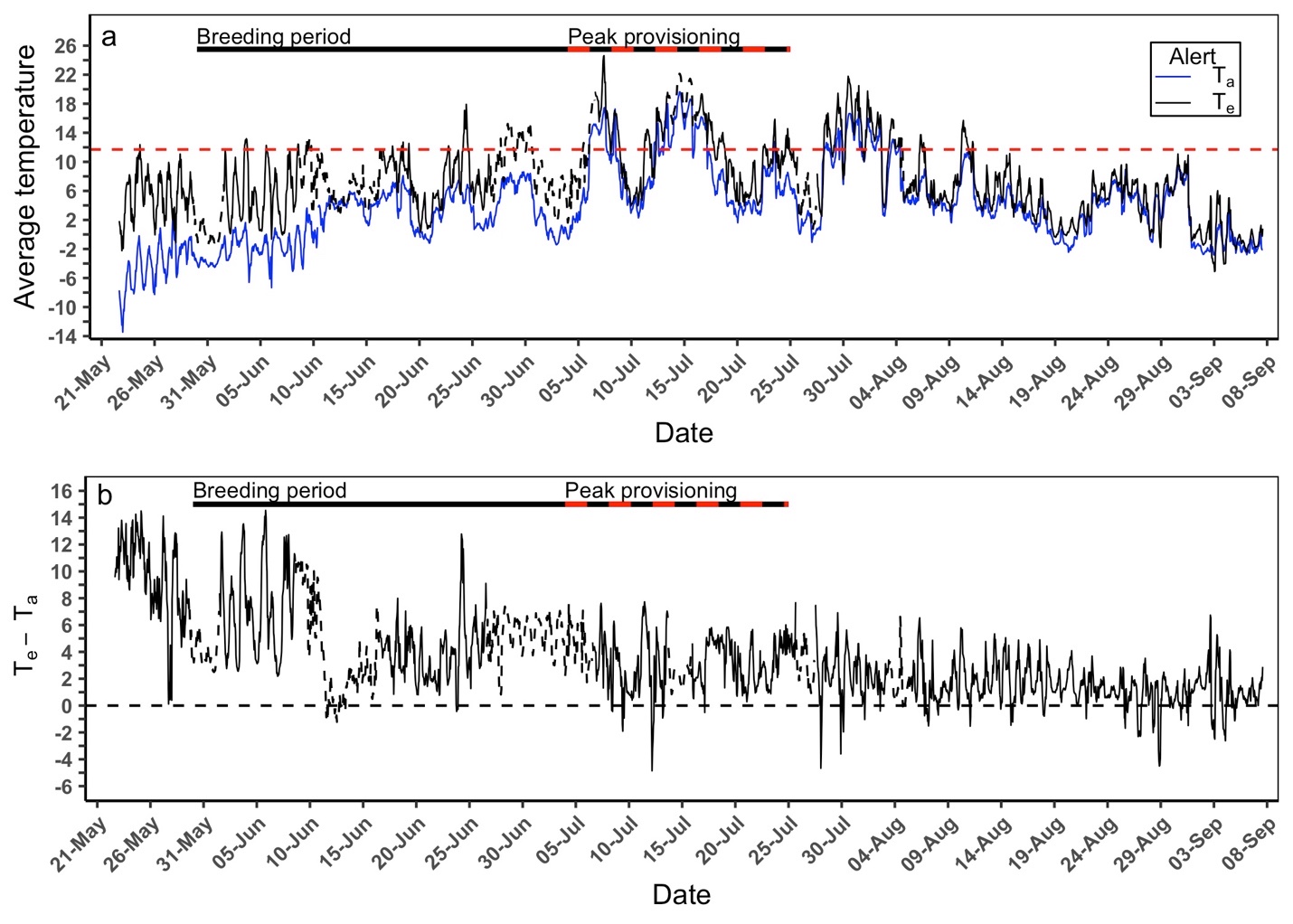

**Figure S1.** (a) Average hourly operative (T_e_) and air (T_a_) temperature and (b) differences between hourly recordings of T_e_ and T_a_ measured from 22 May to 07 September 2019 at Alert, Nunavut Canada. The combined solid black and red/black dashed bar represents the typical breeding period range (30 May to 25 July) while the red/black dashed bar represents the typical peak provisioning period (4 July to 25 July). The horizontal red dashed line in panel (a) represents the 11.7°C operative temperature threshold for birds to maintain thermal balance and sustain an optimal performance ≥ 4x basal metabolic rate. The dashed operative temperature traces represent the predicted operative temperature values from the artificial neural network.

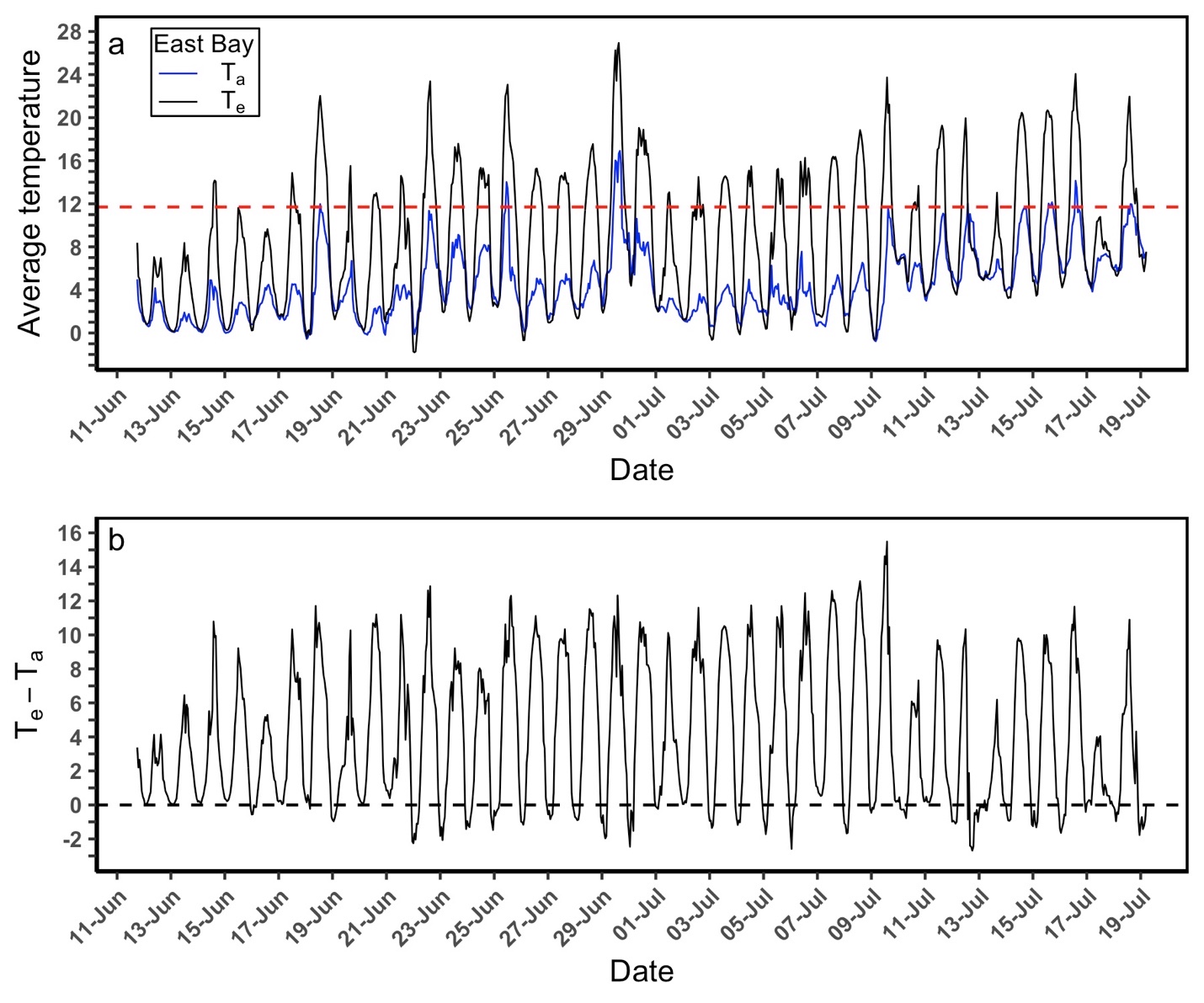

**Figure S2.** (a) Average hourly operative (T_e_) and air (T_a_) temperature values and (b) differences between hourly recordings of T_e_ and T_a_ measured from 11 June to 19 July 2019 at East Bay, Nunavut Canada. The horizontal red dashed line in panel (a) represents the 11.7°C operative temperature threshold for birds to maintain thermal balance and sustain an optimal performance ≥ 4x basal metabolic rate.
